# Human pseudoislet system enables detection of differences in G-protein-coupled-receptor signaling pathways between α and β cells

**DOI:** 10.1101/842989

**Authors:** John T. Walker, Rachana Haliyur, Heather A. Nelson, Matthew Ishahak, Gregory Poffenberger, Radhika Aramandla, Conrad Reihsmann, Joseph R. Luchsinger, Diane C. Saunders, Peng Wang, Adolfo Garcia-Ocaña, Rita Bottino, Ashutosh Agarwal, Alvin C. Powers, Marcela Brissova

## Abstract

G-protein-coupled-receptors (GPCRs) modulate insulin secretion from β cells and glucagon secretion from α cells. Here, we developed an integrated approach to study the function of primary human islet cells using genetically modified pseudoislets that resemble native islets across multiple parameters. We studied the G_i_ and G_q_ GPCR pathways by expressing the designer receptors exclusively activated by designer drugs (DREADDs) hM4Di or hM3Dq. Activation of G_i_ signaling reduced insulin and glucagon secretion, while activation of G_q_ signaling stimulated glucagon secretion but had both stimulatory and inhibitory effects on insulin secretion. Further, we developed a microperifusion system that allowed synchronous acquisition of GCaMP6f biosensor signal and hormone secretory profiles and showed that the dual effects for G_q_ signaling occur through changes in intracellular Ca^2+^. By combining pseudoislets with a microfluidic system, we co-registered intracellular signaling dynamics and hormone secretion and demonstrated differences in GPCR signaling pathways between human β and α cells.

## INTRODUCTION

Pancreatic islets of Langerhans, small collections of specialized endocrine cells interspersed throughout the pancreas, control glucose homeostasis. Islets are composed primarily of β, α, and δ cells but also include supporting cells such as endothelial cells, nerve fibers, and immune cells. Insulin, secreted from the β cells, lowers blood glucose by stimulating glucose uptake in peripheral tissues, while glucagon, secreted from α cells, raises blood glucose through its actions in the liver. Importantly, β and/or α cell dysfunction is a key component of all forms of diabetes mellitus (Brissova et al., 2018; Chen et al., 2017; Cnop et al., 2005; Gloyn et al., 2004; Halban et al., 2014; Haliyur et al., 2018; Hart et al., 2018; Lu and Li, 2018; Naylor et al., 2011; Talchai et al., 2012; Unger and Cherrington, 2012). Thus, an improved understanding of the pathways governing the coordinated hormone secretion in human islets may provide insight into how these may become dysregulated in diabetes.

In β cells, the central pathway of insulin secretion involves glucose entry via glucose transporters where it is metabolized, resulting in an increased ATP:ADP ratio. This shift closes ATP-sensitive potassium channels, depolarizing the cell membrane and opening voltage-dependent calcium channels where calcium influx is a trigger of insulin granule exocytosis (Tokarz et al., 2018). In α cells, the pathway of glucose inhibition of glucagon secretion is not clearly defined with both intrinsic and paracrine mechanisms proposed (Gylfe and Gilon, 2014; Hughes et al., 2018; Yu et al., 2019). Furthermore, gap junctional coupling and paracrine signaling between islet endocrine cells and within the 3D islet architecture are critical for islet function, as individual α or β cells do not show the same coordinated secretion pattern seen in intact islets (Capozzi et al., 2019; Elliott et al., 2014; Svendsen et al., 2018; Unger and Orci, 2010; Zhu et al., 2019).

Although glucose is central to islet hormone secretion, other signals, many of which act through G-protein-coupled-receptors (GPCRs), further modulate and optimize islet hormone secretion (Ahrén, 2009; Persaud, 2017). GPCRs, a broad class of integral membrane proteins, mediate extracellular messages to intracellular signaling through activation of heterotrimeric G-proteins which can be broadly classified into distinct families based on the G_α_ subunit, including G_i_-coupled and G_q_-coupled GPCRs (Weis and Kobilka, 2014). An estimated 30-50% of clinically approved drugs target or signal through GPCRs, including multiple used for treat diabetes treatment (Foord et al., 2005; Hauser et al., 2017). Thus, understanding how GPCR pathways affect human β and α cell secretion is likely to aid in the development of new antidiabetic drugs.

The 3D islet architecture, while essential for function, presents experimental challenges for mechanistic studies of intracellular signaling pathways in primary islet cells. Furthermore, human islets show a number of key differences from rodent islets including their endocrine cell composition and arrangement, glucose set-point, and both basal and stimulated insulin and glucagon secretion, highlighting the importance of studying signaling pathways in primary human cells (Brissova et al., 2005; Cabrera et al., 2006; Dai et al., 2012; 2017; Rodriguez-Diaz et al., 2018).

To study signaling pathways in primary human islet cells within the context of their 3D arrangement, we developed an integrated approach that consists of: 1) human pseudoislets closely mimicking native human islet biology and allowing for efficient genetic manipulation; and 2) a microfluidic system with the synchronous assessment of intracellular signaling dynamics and both insulin and glucagon secretion. Using this system with DREADDs (Armbruster et al., 2007), we demonstrate differences in G_q_ and G_i_ signaling pathways between human β and α cells.

## RESULTS

### Human pseudoislets resemble native human islets and facilitate virally mediated manipulation of human islet cells

To establish an approach that would allow manipulation of human islets, we adapted a system where human islets are dispersed into single cells and then reaggregated into pseudoislets (Arda et al., 2016; Furuyama et al., 2019; Hilderink et al., 2015; Peiris et al., 2018; Yu et al., 2018) (Figure 1A). To optimize the formation and function of human pseudoislets, we investigated two different systems to create pseudoislets, a modified hanging drop system (Foty, 2011; Zuellig et al., 2014) and an ultra-low attachment microwell system. We found both systems generated pseudoislets of comparable quality and function (Figures S1A and S1B) and thus combined groups for comparisons between native islets and pseudoislets. A key determinant of pseudoislet quality was the use of a nutrient- and growth factor-enriched media (termed Vanderbilt pseudoislet media).

**Figure 1.**
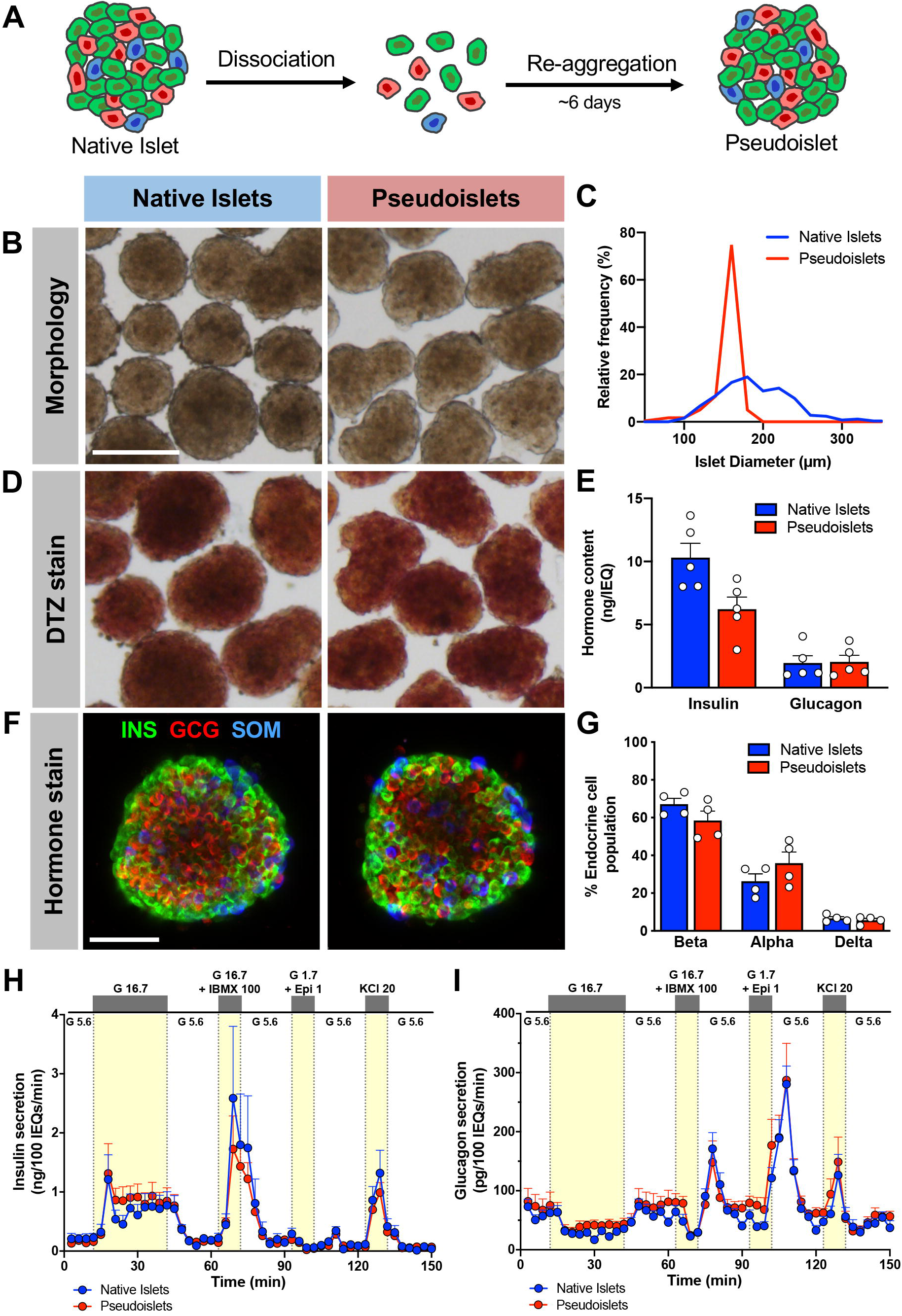
Pseudoislets resemble native human islets in morphology, cell composition, and function. (A) Schematic of pseudoislet formation. (B) Bright-field images showing the morphology of native islets and pseudoislets. Scale bar is 200 μm and also applies to D. (C) Relative frequency plot of islet diameter comparing hand-picked native islets to pseudoislets from the same donor. (D) Dithizone (DTZ) uptake of native islets and pseudoislets. (E) Insulin and glucagon content normalized to islet volume expressed in islet equivalents (IEQs); 1 IEQ corresponds to an islet with a diameter of 150 μm; n=5 donors; p > 0.05. (F) Confocal images of native islets and pseudoislets stained for insulin (INS; β cells), glucagon (GCG; α cells), and somatostatin (SOM; δ cells); scale bar is 50 μm. (G) Quantification of relative endocrine cell composition of native islets and pseudoislets; n=4 donors; p > 0.05. Insulin (H) and glucagon (I) secretory response to various secretagogues measured by perifusion of native islets and pseudoislets from the same donor (n=5). G 5.6 – 5.6 mM glucose; G 16.7 – 16.7 mM glucose; G 16.7 + IBMX 100 – 16.7 mM glucose with 100 μM isobutylmethylxanthine (IBMX); G1.7 + Epi 1 – 1.7 mM glucose and 1μM epinephrine; KCl 20 – 20 mM potassium chloride (KCl). Wilcoxon matched-pairs signed rank test was used to analyze statistical significance in E and G. Panels H and I were analyzed by 2-way ANOVA; p > 0.05. The area under the curve for each secretagogue was compared by one-way ANOVA with Dunn’s multiple comparison test (Figure S1E-S1H and S1J-S1M). Data are represented as mean ± standard error of the mean (SEM).

Pseudoislet morphology, size, and dithizone (DTZ) uptake resembled normal human islets (Figures 1B-1D). Pseudoislet size was controlled to between 150-200 μm in diameter by adjusting the seeding cell density and thus resembled the size of an average native human islet. Compared to native islets from the same donor cultured in parallel using the same pseudoislet media, pseudoislets had similar insulin and glucagon content though insulin content was reduced in pseudoislets from some donors (Figure 1E). Endocrine cell composition was also similar with the ratio of β, α, and δ cells in pseudoislets unchanged compared to cultured native islets from the same donor (Figures 1F and 1G).

As the primary function of the pancreatic islet is to sense glucose and other nutrients and dynamically respond with coordinated hormone secretion, we assessed the function of pseudoislets compared to native islets by perifusion. We used the standard perifusion (termed in the text also as macroperifusion) approach of the Human Islet Phenotyping Program of the Integrated Islet Distribution Program (IIDP; https://iidp.coh.org/) which has assessed nearly 300 human islet preparations. Pseudoislet insulin secretion was indistinguishable from that of native islets in biphasic response to glucose, cAMP-evoked potentiation, epinephrine-mediated inhibition, and KCl-mediated depolarization (Figure 1H). Pseudoislets and native islet also had comparable glucagon secretion, which was inhibited by high glucose, and stimulated by cAMP-mediated processes (IBMX and epinephrine) and depolarization (KCl) (Figure 1I). Compared to native islets on the day of arrival, pseudoislets largely maintained both insulin and glucagon secretion after six days of culture with the exception of a slightly diminished second phase glucose-stimulated insulin secretion (Figure S1C-S1N). These results demonstrate that after dispersion into the single-cell state, human islet cells can reassemble and reestablish intra-islet connections crucial for coordinated hormone release across multiple signaling pathways.

Proliferation, as assessed by Ki67, was low in both native and pseudoislets with β cells below 0.5% and α cells around 1% (Figures 2A and 2B). Similarly, apoptosis, as assessed by TUNEL, was very low (<0.5%) in pseudoislets and cultured human islets (Figures 2A and 2C). Interestingly, the islet architecture of both native whole islets and pseudoislets cultured for six days showed β cells primarily on the islet periphery with α cells and δ cells situated within an interior layer. Furthermore, the core of both the cultured native islets and pseudoislets consisted largely of extracellular matrix (collagen IV) and endothelial cells (caveolin-1) (Figures 2A, 2D, and 2E), likely reflective of the consequences of culture. The survival of intraislet endothelial cells in culture for an extended period of time could be due to the nutrient- and growth factor-enriched media. Additionally, the islet cell arrangement suggests that extracellular matrix and endothelial cells may facilitate pseudoislet assembly.

**Figure 2.**
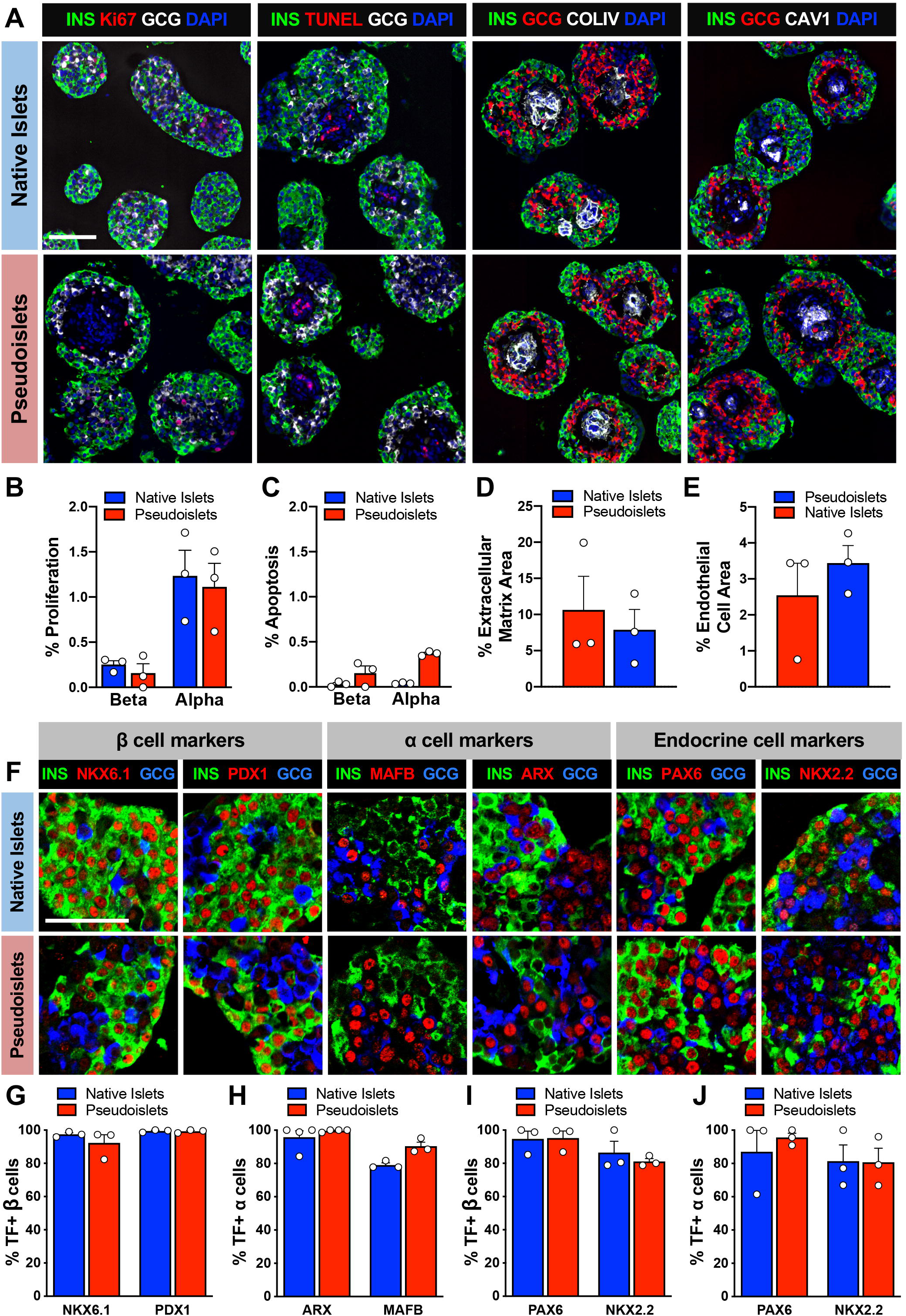
Pseudoislets resemble native human islets in proliferation, apoptosis, architecture, and express markers of α and β cell identity. (A) Immunofluorescence visualization of labeling for insulin (INS; β cells) and (GCG; α cells) in combination with detection of proliferation (Ki67), apoptosis (TUNEL), extracellular matrix (collagen IV, COLIV) and endothelial cells (caveolin-1, CAV1). Scale bar is 100 µm. (B) Quantification of β and α cell proliferation in native islets and pseudoislets; expressed as % INS+ or GCG+ cells expressing Ki67; n=3 donors; p > 0.05. (C) Quantification of β and α cell apoptosis by TUNEL assay; n=3 donors; p > 0.05. (D) Quantification of COLIV-expressing extracellular matrix; expressed as % COLIV+ area to INS+ and GCG+ cell area; n=3 donors; p > 0.05. (E) Quantification of endothelial cell area; expressed as % CAV1+ cell area to INS+ and GCG+ cell area; n=3 donors; p > 0.05. (F) Expression of transcription factors (TF) important for β cell identity (NKX6.1 and PDX1), α cell identity (MAFB and ARX), and pan endocrine cell identity (PAX6 and NKX2.2). Scale bar is 50 µm. (G) Quantification β cell identity markers in β cells of native islets and pseudoislets (n=3 donors/marker; p > 0.05). (H) Quantification of α cell identity markers in α cells of native islets and pseudoislets (n=3-4 donors/marker; p > 0.05). (I-J) Quantification of pan endocrine markers in β (I) and α (J) cells of native islets and pseudoislets (n=3 donors/marker; p > 0.05). Wilcoxon matched-pairs signed rank test was used to analyze statistical significance in panels B-E and G-J. Data are represented as mean ± SEM.

To assess markers of α and β cell identity in pseudoislets, we investigated expression of several key islet-enriched transcription factors. The expression of β (PDX1, NKX6.1) and α cell markers (MAFB, ARX) as well as those expressed in both cell types (PAX6, NKX2.2) was maintained in pseudoislets when compared to native islets (Figures 2F-2J), indicating that the process of dispersion and reaggregation does not affect islet cell identity.

The 3D structure of intact islets makes virally mediated manipulation of human islet cells challenging due to poor viral penetration into the center of the islet. We adopted the pseudoislet system to overcome this challenge by transducing the dispersed single islet cells before reaggregation (Figure 3A). To optimize transduction efficiency and subsequent pseudoislet formation, we incubated with adenovirus for 2 hours in Vanderbilt pseudoislet media at a multiplicity of infection (MOI) of 500. Transducing pseudoislets with control adenovirus did not affect pseudoislet morphology or function and achieved high transduction efficiency of β and α cells throughout the entire pseudoislet (Figures S2A-S2E). Interestingly, β cells showed a higher transduction efficiency (90%) than α cells (70%), suggesting that α cells may be inherently more difficult to transduce with adenovirus (Figure S3B).

**Figure 3.**
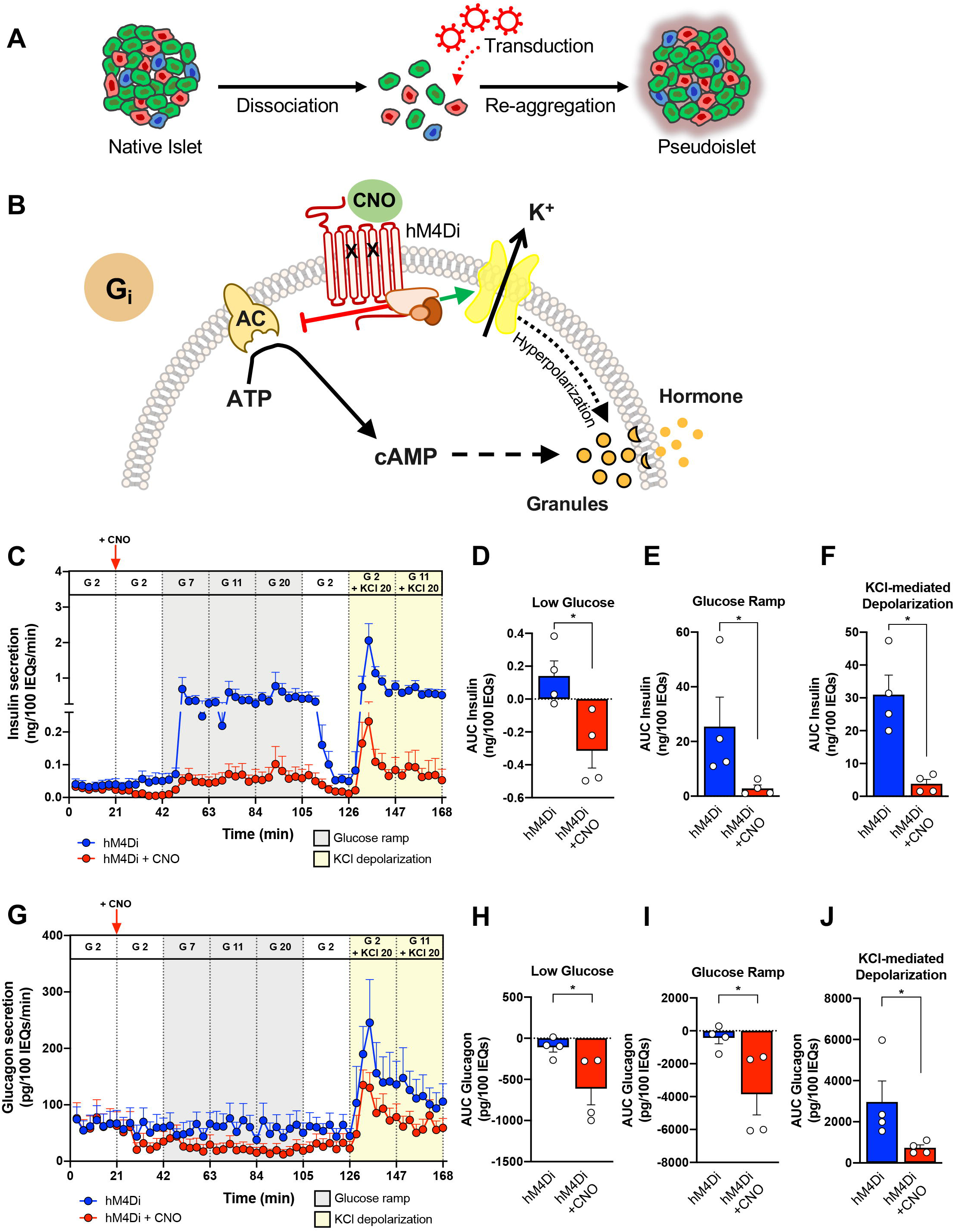
G_i_ activation reduces insulin and glucagon secretion. (A) Schematic of incorporation of efficient viral transduction into pseudoislet approach. (B) Schematic of the G_i_-coupled GPCR signaling pathway. CNO – clozapine-N-oxide, AC – adenylyl cyclase, ATP –adenosine triphosphate, cAMP – cyclic adenosine monophosphate, K^+^ – potassium ion. (C) Dynamic insulin secretion assessed by macroperifusion in response to low glucose (G 2 – 2 mM glucose; white), glucose ramp (G 7 – 7 mM, G 11 – 11 mM, and G 20 – 20 mM glucose; grey) and KCl-mediated depolarization (KCl 20 – 20 mM potassium chloride in the presence of G 2 or G 11; yellow) in the absence (blue trace) or presence of CNO (red trace); n=4 donors/each. 10 µM CNO was added after the first period of 2 mM glucose as indicated by a vertical red arrow and then continuously administered for the duration of the experiment (red trace). Note the split of y-axis to visualize differences between traces at G 2 ± CNO. (D-F) Insulin secretion was integrated by calculating the area under the curve (AUC) for response to the low glucose (white), glucose ramp (gray), and KCl-mediated depolarization (yellow). Baseline was set to the average value of each trace from 0 to 21 minutes (before CNO addition). (G-J) Glucagon secretion was analyzed in parallel with insulin as described above. Insulin and glucagon secretory traces in panels C and G, respectively, were compared in the absence vs. presence of CNO by two-way ANOVA; ****, p < 0.0001 for both insulin and glucagon secretion. Area under the curve of insulin (D-F) and glucagon responses (H-J) to low glucose, glucose ramp, and KCl-mediated depolarization were compared in the absence vs. presence of CNO by Mann-Whitney test; *, p < 0.05. Data are represented as mean ± SEM.

### Activation of G_i_ signaling reduces insulin and glucagon secretion

Studying GPCR signaling with endogenous receptors and ligands can be complicated by a lack of specificity—ligands that can activate multiple receptors or receptors that can be activated by multiple ligands. To overcome these limitations, we employed the DREADD technology. DREADDs are GPCRs with specific point mutations that render them unresponsive to their endogenous ligand. Instead, they can be selectively activated by the otherwise inert ligand, clozapine-N-oxide (CNO), thus providing a selective and inducible model of GPCR signaling (Armbruster et al., 2007; Wess, 2016). DREADDs are commonly used in neuroscience as molecular switches to activate or repress neurons with G_q_ or G_i_ signaling, respectively (Roth, 2016). In contrast, there have been comparatively very few studies using DREADDs in the field of metabolism, but they have included investigating the G_q_ DREADD in mouse β cells and the G_i_ DREADD in mouse α cells (Guettier et al., 2009; Zhu et al., 2019). To our knowledge, this is the first study to utilize this powerful technology in human islets.

To investigate G_i_-coupled GPCR signaling, we introduced adenovirus encoding hM4Di (Ad-CMV-hM4Di-mCherry), a G_i_ DREADD, into dispersed human islet cells, allowed reaggregation into pseudoislets and then tested the effect of activated G_i_ signaling (Figure 3A). G_i_-coupled GPCRs signal by inhibiting adenylyl cyclase, thus reducing cAMP, and by activating inward rectifying potassium channels (Figure 3B). CNO had no effect on insulin or glucagon secretion in mCherry-expressing pseudoislets (Figures S2F and S2G), thus we compared the dynamic hormone secretion of hM4Di-expressing pseudoislets with and without CNO in response to multiple stimuli by perifusion. Activation of G_i_ signaling had clear inhibitory effects on insulin secretion by β cells at low glucose, which became more prominent with progressively higher glucose concentrations (gray shading; Figures 3C-3E). Furthermore, bypassing glucose metabolism by directly activating β cells via depolarization with potassium chloride did not overcome this inhibition by G_i_ signaling (yellow shading; Figures 3C and 3F). Together these data demonstrate that in human β cells, G_i_ signaling significantly attenuates, but does not completely prevent, insulin secretion and that this effect, at least in part, occurs downstream of glucose metabolism.

The activation of G_i_ signaling had also inhibitory effects on glucagon secretion (Figures 3G-3J). When stimulated with potassium chloride, pseudoislets with activated G_i_ signaling increased glucagon secretion but not to the level of controls. This demonstrates that the inhibitory effects of G_i_ signaling persist even if the α cell is directly activated by depolarization. Thus, in α cells, activation of G_i_ signaling reduces glucagon secretion across a range of glucose levels and when the cell is depolarized by potassium chloride.

### Activation of G_q_ signaling greatly stimulates glucagon secretion but has both stimulatory and inhibitory effects on insulin secretion

G_q_-coupled GPCRs signal through phospholipase C, leading to IP_3_-mediated Ca^2+^ release from the endoplasmic reticulum (Figure 4A). To investigate G_q_-coupled GPCR signaling, we introduced hM3Dq (Ad-CMV-hM3Dq-mCherry), a G_q_ DREADD, into dispersed human islet cells, allowed reaggregation, and assessed hM3Dq-expressing pseudoislets by perifusion. When CNO was added to activate G_q_ signaling, there was an acute increase in insulin secretion. However, this was not sustained as insulin secretion fell quickly back to baseline, highlighting a dynamic response to G_q_ signaling in β cells (Figures 4B-4E). Furthermore, continued G_q_ activation inhibited glucose-stimulated insulin secretion, suggesting that in certain scenarios, G_q_ signaling may override the ability of glucose to stimulate insulin secretion. These results highlight the value of assessing hormone secretion in the dynamic perifusion system. Finally, G_q_ activation reduced, but did not completely prevent, insulin secretion in response to direct depolarization with potassium chloride, indicating that the inhibitory effects cannot be overcome by bypassing glucose metabolism and suggesting that they occur downstream of the K_ATP_ channel. Together, these data indicate that activated G_q_ signaling can have both stimulatory and inhibitory effects on human β cells.

**Figure 4.**
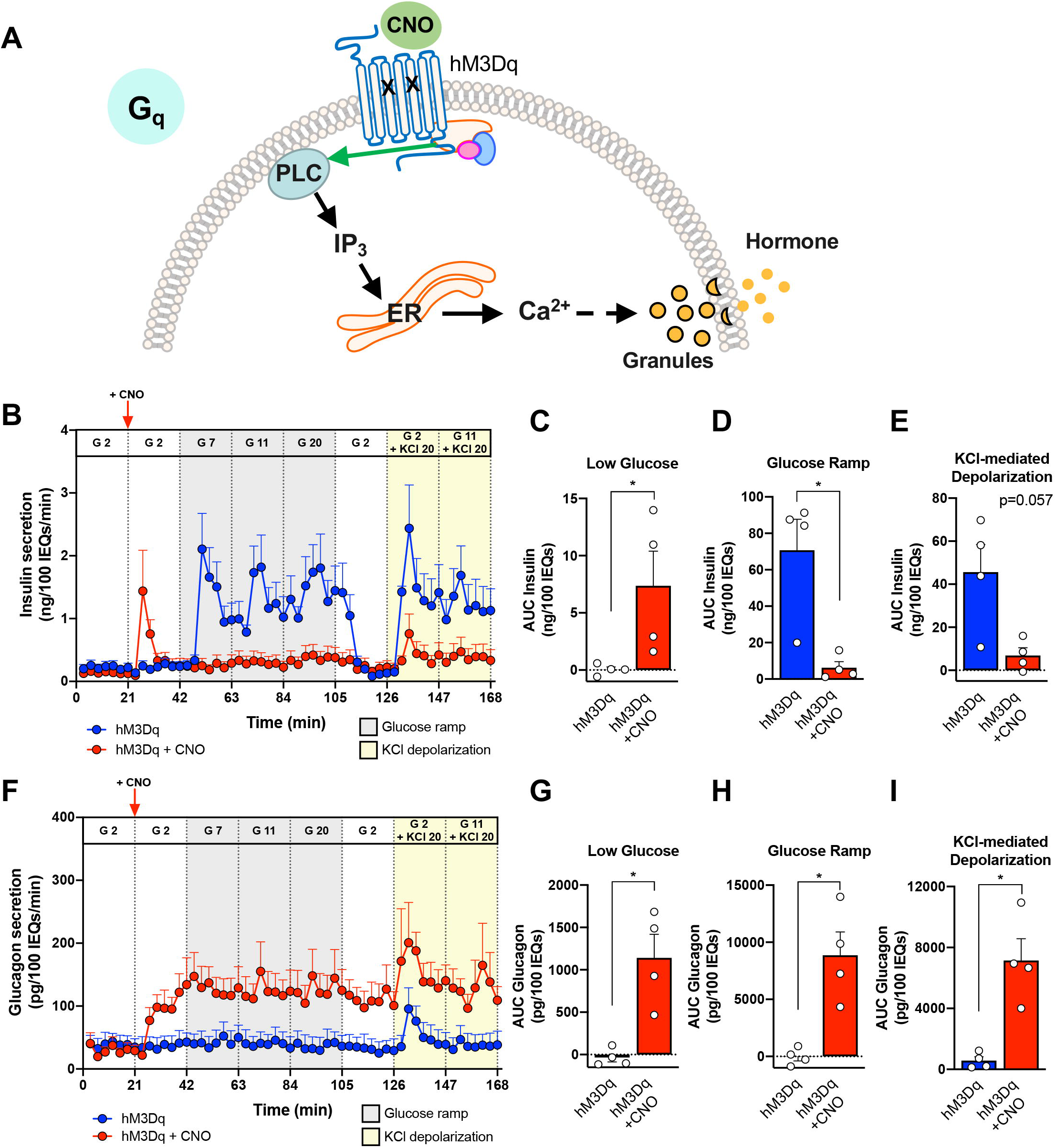
G_q_ activation stimulates glucagon secretion but has stimulatory and inhibitory effects on insulin secretion. (A) Schematic of the G_q_-coupled GPCR signaling pathway. CNO – clozapine-N-oxide, PLC – phospholipase C, IP_3_ – inositol triphosphate, ER – endoplasmic reticulum, Ca^2+^ – calcium ion. (B-E) Dynamic insulin secretion was assessed by macroperifusion and analyzed as described in detail in Figure 3; n=4 donors/each. (F-I) Glucagon secretion was analyzed in parallel with insulin as described in Figure 3. Insulin and glucagon secretory traces in panels B and F, respectively were compared in the absence vs. presence of CNO by two-way ANOVA; ****, p < 0.0001 for both insulin and glucagon secretion. Area under the curve of insulin (C-E) and glucagon responses (G-I) to each stimulus were compared in the absence vs. presence of CNO by Mann-Whitney test; *, p < 0.05. Data are represented as mean ± SEM.

In contrast, activation of G_q_ signaling in α cells robustly increased glucagon secretion in low glucose and it remained elevated with continued CNO exposure during glucose ramp as well as in the presence of potassium chloride (Figures 4F-4I). This indicates that in contrast to the β cells, activation of G_q_ signaling in the α cells robustly stimulates glucagon secretion and this response is sustained across a glucose ramp and during KCl-mediated depolarization.

### Integration of pseudoislet system with genetically-encoded biosensor and microfluidic device allows synchronous measurement of intracellular signals and hormone secretion

While conventional macroperifusion systems, including the perifusion system used in this study reliably assess islet hormone secretory profiles (Brissova et al., 2018; Haliyur et al., 2018; Hart et al., 2018; Kayton et al., 2015; Saunders et al., 2019), their configuration does not allow coupling with imaging systems to measure intracellular signaling. To overcome this challenge, we developed an integrated microperifusion system consisting of pseudoislets and a microfluidic device that enables studies of islet intracellular signaling using genetically-encoded biosensors in conjunction with hormone secretion (Figures 5A and S3A-S3C). The microfluidic device (Figure S3A) (Lenguito et al., 2017) is made of bio-inert and non-absorbent materials with optimized design for nutrient delivery, synchronous islet imaging by confocal microscopy, and collection of effluent fractions for analysis of insulin and glucagon secretion. The microperifusion system uses smaller volumes, slower flow rates, and fewer islets than our conventional macroperifusion system (Figures S3D-S3F).

**Figure 5.**
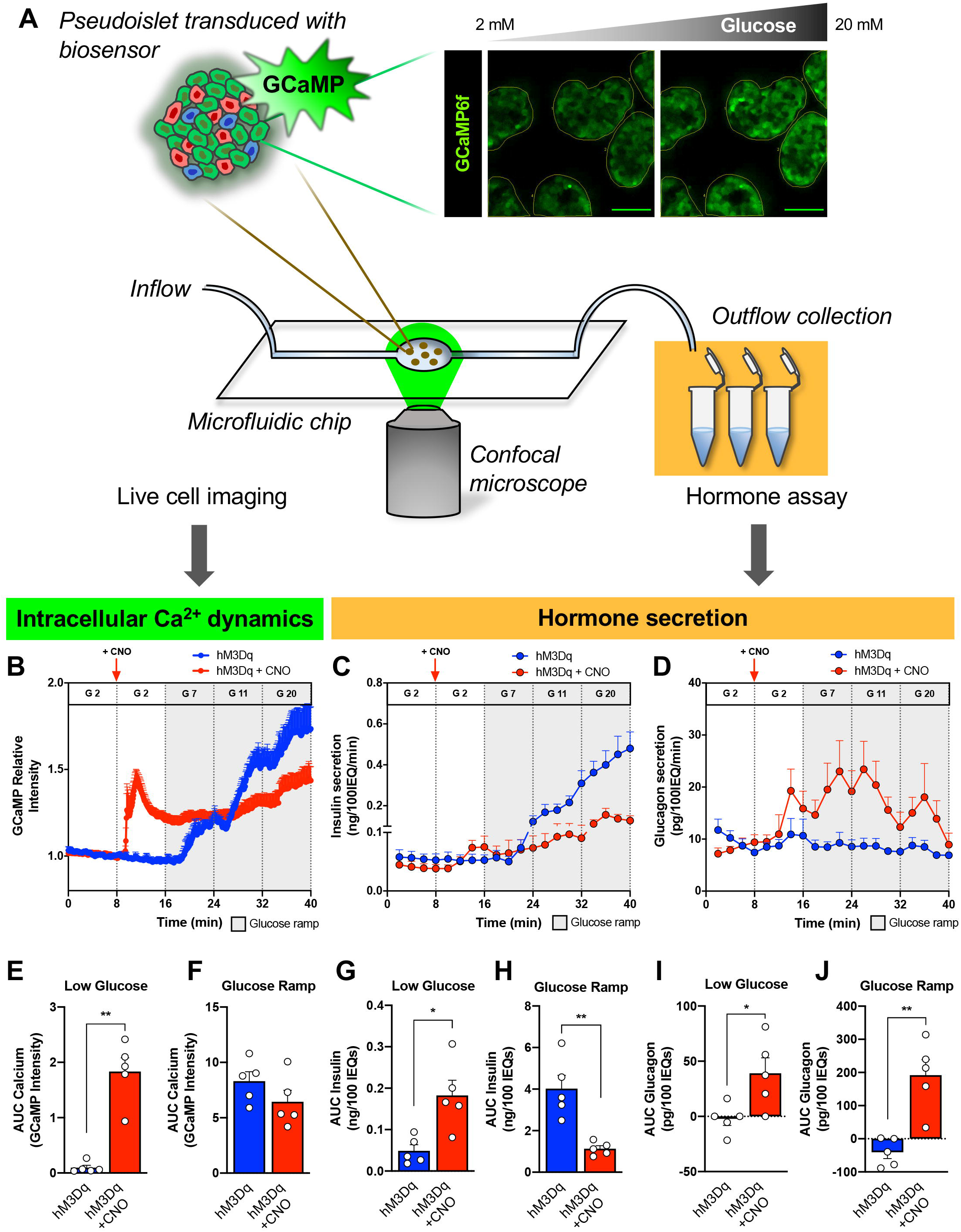
Pseudoislet system integrated with microfluidic device allows for co-registration of hormone secretion and intracellular signaling dynamics. (A) Schematic of pseudoislet system integration with a microfluidic device to allow for synchronous detection of intracellular signaling dynamics by the genetically encoded GCaMP6f biosensor and confocal microscopy, and collection of microperifusion efflux for hormone analysis. Dynamic changes in GCaMP6f relative intensity (B), insulin secretion (C), and glucagon secretion (D) assessed during microperifusion in response to a low glucose (G 2 – 2 mM glucose; white), glucose ramp (G 7 – 7 mM, G 11 – 11 mM, and G 20 – 20 mM glucose; grey) and in the absence (blue trace) or presence of CNO (red trace); n=3 donors/each. 10 µM CNO was added after the first period of 2 mM glucose as indicated by a vertical red arrow and then continuously administered for the duration of the experiment (red trace). See Supplemental Videos 1 and 2 for representative visualization of each experiment. Calcium signal (E, F) and insulin (G, H) and glucagon (I, J) secretion was integrated by calculating the area under the curve (AUC) for response to the low glucose (white) and glucose ramp (gray). Baseline was set to the average value of each trace from 0 to 8 minutes (before CNO addition). Calcium and hormone traces in panels B-D were compared in the absence vs. presence of CNO by two-way ANOVA; * p < 0.05 for calcium trace, **** p < 0.0001 for both insulin and glucagon secretion. Area under the curve of calcium (E, F), insulin (G, H) and glucagon responses (I, J) to low glucose and glucose ramp were compared in the absence vs. presence of CNO by Mann-Whitney test; *, p < 0.05, **, p < 0.01. Data are represented as mean ± SEM.

To investigate the dual effects of activated G_q_ signaling on insulin secretion, we co-transduced pseudoislets with hM3Dq and GCaMP6f (Figure S3C), a calcium biosensor (Ad-CMV-GCaMP6f), as the G_q_ pathway conventionally signals through intracellular Ca^2+^ (Figure 4A). In the absence of CNO, hM3Dq-expressing pseudoislets had stepwise increases in GCaMP6f relative intensity as glucose increased, corresponding to increasing intracellular Ca^2+^ and highlighting the added value of the system (Figure 5B). This intracellular Ca^2+^ response to stepwise glucose increase was accompanied by increasing insulin secretion (Figure 5C), but the first phase of insulin secretion was not as clearly resolved as in the macroperifusion (Figure 4B). Since the design of the macro- and microperifusion systems is significantly different, we used multiphysics computational modeling with finite element analysis (Buchwald, 2011; Buchwald et al., 2018) to model the insulin secretion dynamics of the two systems (Figures S3H and S3I). This modeling accurately predicted the overall shape of each insulin secretory trace with the macroperifusion showing a “saw-tooth” pattern (Figure S3H) while the microperifusion had a more progressive increase (Figure S3I). Using this approach, we found that differences in the insulin secretory profiles were primarily due to the different fluid dynamics and experimental parameters between the two perifusion systems, especially the experimental time for each stimulus and the flow rate. Overall, this analysis demonstrates how perifusion parameters can impact insulin secretory pattern and indicates the strength of using complementary approaches.

When G_q_ signaling was activated with CNO, we again saw a transient stimulation of insulin secretion at low glucose followed by relative inhibition through the glucose ramp, while glucagon secretion from α cells was stimulated throughout the entire perifusion, independently of glucose concentration (Figures 5C, 5D, 5G-5J). Furthermore, the Ca^2+^ dynamics in response to G_q_ activation were consistent with the insulin secretory trace showing a rapid but short-lived increase in intracellular Ca^2+^. Interestingly, the Ca^2+^ signal remained elevated above baseline but did not significantly increase with rising glucose (Figures 5B,5E and 5F). This indicates that the dual effects of G_q_ signaling on insulin secretion in β cells are largely mediated by changes in intracellular Ca^2+^ levels.

## DISCUSSION

The three-dimensional multicellular human islet architecture, while essential for islet cell function presents experimental challenges for mechanistic studies of intracellular signaling pathways. Using primary human islets, we developed a pseudoislet system that resembles native human islets in morphology, cellular composition, cell identity, and dynamic insulin and glucagon secretion. This system allows for efficient virally mediated genetic manipulation in almost all cells in the pseudoislet. To evaluate the coordination between intracellular signals and islet hormone secretion, we developed an integrated system consisting of pseudoislets and a microfluidic device that enables studies of islet intracellular signaling using genetically encoded biosensors in conjunction with hormone secretion. Furthermore, we used this integrated approach to define new aspects of human islet biology by investigating GPCR signaling pathways using DREADDs and a calcium biosensor.

Despite α and β cells both being excitable secretory cells and sharing many common developmental and signaling components, they show similar and distinct responses to activation of GPCR signaling pathways, highlighting the uniqueness in each cell’s molecular machinery. The activation of G_i_ signaling was inhibitory in both β and α cells resulting in reduced insulin and glucagon secretion, respectively, and showed more substantial impact in β cells where this signaling blunted insulin response to both a glucose ramp and to KCl-mediated depolarization. Interestingly, direct KCl depolarization was not sufficient to overcome these inhibitory effects in either cell type, suggesting that reduced cAMP via the inhibition of adenylyl cyclase plays an important role in both insulin and glucagon secretion. These results align well with recent studies in β cells suggesting cAMP tone is crucial for insulin secretion and observations in α cells highlighting cAMP as a key mediator of glucagon secretion (Capozzi et al., 2019; Elliott et al., 2014; Tengholm and Gylfe, 2017; Yu et al., 2019).

The activation of G_q_ signaling showed major differences in β and α cells. In α cells, the activated G_q_ signaling elicited a robust and sustained increase in glucagon secretion in the presence of a glucose ramp and potassium chloride. In contrast, G_q_ signaling in β cells had a transient stimulatory effect in low glucose and then inhibitory effects on both insulin and intracellular Ca^2+^ levels with sustained activation during glucose ramp. Interestingly, previous studies of acetylcholine signaling have also reported dual effects on Ca^2+^ dynamics in β cells depending on the length of stimulation (Gilon et al., 1995). This signaling was thought to be mediated through the muscarinic acetylcholine receptor M_3_ (from which the hM3Dq DREADD is based). Overall, these results suggest a negative feedback or protective mechanism that prevents sustained insulin release from β cells in response to G_q_ signaling that is not active in α cells under similar circumstances.

There are limitations and caveats to the current study. First, our approach expressed the DREADD receptors in all cell types. Although we can distinguish the effects on β and α cells through the cell’s distinct hormone, it is possible that paracrine signaling, including somatostatin from δ cells, is contributing to the results described here. Future modifications could incorporate cell-specific promoters to target a particular islet cell type. Second, the DREADD receptors are likely expressed at higher levels than endogenous GPCRs. To mitigate this, we used the appropriate DREADD-expressing pseudoislets as our controls and were encouraged to see normal secretory responses in these control pseudoislets. Third, while there is some concern that CNO can be reverse-metabolized in vivo into clozapine which could potentially have off-target effects (Gomez et al., 2017), this is unlikely in our in vitro system. We also verified that CNO had no effect on mCherry-expressing pseudoislets. Fourth, we used a CNO concentration of 10 µM for all of our experiments, a standard concentration used for in vitro assays (Smith et al., 2016; Zhu et al., 2019), but it is possible that islet cells may show dose-dependent effects. Finally, this is an in vitro study, and there may be differences in these pathways compared to what is seen in vivo. Future work could involve transplantation of DREADD-expressing pseudoislets into immunodeficient mice to study the effect of activating these pathways on human islets in vivo (Dai et al., 2017).

Overall, these findings demonstrate the utility of the pseudoislet system for the ability to manipulate human islets. Other approaches include inducible pluripotent stem cells that allow similar genetic manipulation. However, it is unclear if these approaches create normal human islet cells. We show in this system that α and β cells maintain their fully differentiated state as well as their dynamic responsiveness to glucose and other stimuli. Additionally, this approach allows for the study of all islet cells within the context of other cell types and 3D assembly.

Here, we focus on virally mediated gene expression to alter signaling pathways, but the system could be adapted to accommodate technologies such as CRISPR. Furthermore, after islet dispersion into single cells, techniques to purify live cell populations such as fluorescence-activated cell sorting (FACS) with cell surface antibodies (Dorrell et al., 2008; Saunders et al., 2019) could be incorporated to allow manipulation of the pseudoislet cellular composition as well as cell-specific gene manipulation. Ultimately, the integration of the pseudoislet approach with a microfluidic perifusion system and live cell imaging provides a powerful experimental platform to gain insight into human islet biology and the mechanisms controlling regulated islet hormone secretion.

## ACKNOWLEDGMENTS

We thank the organ donors and their families for their invaluable donation and the International Institute for Advancement of Medicine (IIAM), Organ Procurement Organizations, National Disease Research Exchange (NDRI), and the Alberta Diabetes Institute IsletCore for their partnership in studies of human pancreatic tissue for research. This study used human pancreatic islets that were provided by the NIDDK-funded Integrated Islet Distribution Program at the City of Hope (NIH Grant # 2UC4 DK098085). Experiments were performed in part through the use of the Vanderbilt Cell Imaging Shared Resource (supported by NIH grants CA68485, DK20593, DK58404, DK59637 and EY08126). This research was performed using resources and/or funding provided by the National Institute of Diabetes and Digestive and Kidney Diseases–supported HIRN (RRID:SCR_014393; https://hirnetwork.org; UC4 DK104211, DK108120, DK112232, and DK120456), by DK106755, DK117147, DK94199, T32GM007347, F30DK118830, F31DK118860, P30DK020541, and DK20593, and by grants from JDRF (2-SRA-2019-699-S-B), The Leona M. and Harry B. Helmsley Charitable Trust, and the Department of Veterans Affairs (BX000666).

## AUTHOR CONTRIBUTIONS

JTW, RH, HAN, MI, JRL, AA, MB, ACP conceived and designed the experiments. JTW, RH, HAN, GP, RA, CR, DCS, MI, PW, AGO, RB, and MB performed experiments or analyzed the data and interpreted results. JTW, MB, and ACP wrote the manuscript. All authors reviewed, edited, and approved the final version.

## METHODS

### Human islet isolation

Human islets (n=24 preparations, Table S1) were obtained through partnerships with the Integrated Islet Distribution Program (IIDP, http://iidp.coh.org/), Alberta Diabetes Institute (ADI) IsletCore (https://www.epicore.ualberta.ca/IsletCore/), Human Pancreas Analysis Program (https://hpap.pmacs.upenn.edu/), or isolated at the Institute of Cellular Therapeutics of the Allegheny Health Network (Pittsburgh, PA). Assessment of human islet function was performed by islet macroperifusion assay on the day of arrival, as previously described (Kayton et al., 2015). Primary human islets were cultured in CMRL 1066 media (5.5 mM glucose, 10% FBS, 1% Pen/Strep, 2 mM L-glutamine) in 5% CO_2_ at 37°C for <24 hours prior to beginning studies. The Vanderbilt University Institutional Review Board does not classify de-identified human pancreatic specimens as human subject research.

This study used data from the Organ Procurement and Transplantation Network (OPTN) that was in part compiled from the Data Hub accessible to IIDP-affiliated investigators through IIDP portal (https://iidp.coh.org/secure/isletavail). The OPTN data system includes data on all donors, wait-listed candidates, and transplant recipients in the US, submitted by the members of the Organ Procurement and Transplantation Network (OPTN). The Health Resources and Services Administration (HRSA), U.S. Department of Health and Human Services provides oversight to the activities of the OPTN contractor. The data reported here have been supplied by UNOS as the contractor for the Organ Procurement and Transplantation Network (OPTN). The interpretation and reporting of these data are the responsibility of the author(s) and in no way should be seen as an official policy of or interpretation by the OPTN or the U.S. Government.

### Pseudoislet formation

Briefly, human islets were handpicked to purity and then dispersed with HyClone trypsin (Thermo Scientific). Islet cells were counted and then seeded at 2000 cells per well in CellCarrier Spheroid Ultra-low attachment microplates (PerkinElmer) or 2500 cells per drop in GravityPLUS™ Hanging Drop System (InSphero) in enriched Vanderbilt pseudoislet media. Cells were allowed to reaggregate for 6 days before being harvested and studied.

### Immunohistochemical Analysis

Immunohistochemical analysis of islets was performed by whole-mount or on 8-μm cryosections of islets embedded in collagen gels as previously described (Brissova et al., 2005; 2018). Primary antibodies to all antigens and their working dilutions are listed in Table S2. Apoptosis was assessed by TUNEL (Millipore, S7165) following the manufacturer’s instructions. Digital images were acquired with a Zeiss LSM 880 or LSM 510 laser-scanning confocal microscope (Zeiss Microscopy Ltd, Jena, Germany) or ScanScope FL (Aperio/Leica Biosystems, Wetzler, Germany). Images were analyzed using HALO Image Analysis Software (Indica Labs, Albuquerque, New Mexico) or MetaMorph v7.1 (Molecular Devices LLC, San Jose, CA).

### Adenovirus

Adenoviral vectors CMV-mCherry (VB180905-1046uck), CMV-hM4Di-mCherry (VB180904-1144bbp), CMV-hM3Dq-mCherry (VB160707-1172csx) were constructed by VectorBuilder Inc (Chicago, IL) and adenovirus was prepared, amplified, and purified by the Human Islet and Adenovirus Core of the Einstein-Sinai Diabetes Research Center (New York, NY) or Welgen Inc (Worcester, MA). Titers were determined by plaque assay. Ad-CMV-GCaMP6f was purchased from Vector Biolabs (Catalog #1910, Malvern, PA). Dispersed human islets were incubated with adenovirus at a multiplicity of infection of 500 for 2 hours in Vanderbilt pseudoislet media before being spun, washed, and plated.

### Assessment of islet function in vitro by static incubation

Pseudoislets (10-20 IEQs/well) were placed in 2 mL of DMEM (media, 2mM glucose) of a 12-well plate (351143, Corning) and allowed to equilibrate for 30 minutes and then were transferred to media containing the stimuli of interest for 40 minutes. Media from this incubation was assessed for insulin and glucagon by radioimmunoassay (insulin: RI-13K, glucagon: GL-32K, Millipore) as previously reported (Brissova et al., 2018).

### Assessment of islet function by macroperifusion

Function of native islets and pseudoislets was studied in a dynamic cell perifusion system at a perifusate flow rate of 1 mL/min as described in (Brissova et al., 2018; Kayton et al., 2015) using approximately 250 IEQs/chamber. The effluent was collected at 3-minute intervals using an automatic fraction collector. Insulin and glucagon concentrations in each perifusion fraction and islet extracts were measured by radioimmunoassay (insulin: RI-13K, glucagon: GL-32K, Millipore, Burlington, MA).

### Microperifusion platform

The microperifusion platform (Figures 5 and S3) is based on a previously published microfluidic device with modifications (Lenguito et al., 2017). Design modifications were incorporated using SolidWorks 2018 3D computer-aided design (CAD) software. Microfluidic devices were machined, according to the CAD models, using a computer numerical controlled milling machine (MDX-540, Roland) from poly(methyl methacrylate) workpieces. To reduce the optical working distance, through-holes were milled into the culture wells and a #1.5 glass coverslip was bonded to the bottom component of the microfluidic device using a silicone adhesive (7615A21, McMaster-Carr). Custom gaskets were fabricated using a two-part silicone epoxy (Duraseal 1533, Cotronics Corp) and bonded into the top component of the device using a specialized polyester adhesive (PS-1340, Polymer Science). The two components of the microfluidic device (Figure S3A) are assembled in a commercially available device holder (Fluidic Connect PRO with 4515 Inserts, Micronit Microfluidics), which creates a sealed system and introduces fluidic connections to a peristaltic pump (Instech, P720) though 0.01” FEP tubing (IDEX, 1527L) and a low volume bubble trap (Omnifit, 006BT) placed in the fluid line just before the device inlet to prevent bubbles from entering the system (see Figure S3B for microperifusion assembly).

### Assessment of pseudoislets by microperifusion

The microperifusion apparatus was contained in a temperature-controlled incubator (37°C) fitted to a Zeiss LSM 880 laser-scanning confocal microscope (Zeiss Microscopy Ltd, Jena, Germany) (Figure S3B). Pseudoislets (approximately 25 IEQs/chamber) were loaded in a pre-wetted well, imaged with a stereomicroscope to determine loaded IEQ, and perifused at 100 μL/min flow rate with Krebs-Ringer buffer containing 125 mM NaCl, 5.9 mM KCl, 2.56 mM CaCl_2_, 1 mM MgCl_2_, 25 mM HEPES, 0.1% BSA, pH 7.4 at 37°C. Perifusion fractions were collected at 2-minute intervals following a 20-minute equilibration period in 2 mM glucose using a fraction collector (Bio-Rad, Model 2110) and analyzed for insulin and glucagon concentration by RIA (insulin –RI-13K, glucagon – GL-32K, Millipore). GCaMP6f biosensor was excited at 488 nm and fluorescence emission detected at 493 – 574 nm. Images were acquired at 15-μm depth every 5 seconds using a 20x/0.80 Plan-Apochromat objective. Image analysis was performed with MetaMorph v7.1 software (Molecular Devices, San Jose, CA). Pseudoislets in the field of view (3-7 pseudoislets/field) were annotated using the region of interest tool. The GCaMP6f fluorescence intensity recorded for each time point was measured across annotated pseudoislet regions and normalized to the baseline fluorescence intensity acquired over the 60 seconds in 2 mM glucose prior to stimulation. The calcium, insulin, and glucagon traces were averaged from 5 microperifusion experiments from 3 independent donors.

### Fluid dynamics and mass transport modeling

Two-dimensional (2D) finite element method (FEM) models, which incorporate fluid dynamics, mass transport, and islet physiology, were developed for the macroperifusion and microperifusion platforms and implemented in COMSOL Multiphysics Modeling Software (Release Version 5.0). Fluid dynamics were governed by the Navier-Stokes equation for incompressible Newtonian fluid flow. Convective and diffusive transport of oxygen, glucose, and insulin were governed by the generic equation for transport of a diluted species in the chemical species transport module. Islet physiology was based on Hill (generalized Michaelis-Menten) kinetics using local concentrations of glucose and oxygen, as previously described (Buchwald, 2011; Buchwald et al., 2018). The geometry of the macroperifusion platform was modeled as the 2D cross-section of a cylindrical tube with fluid flowing from bottom to top (Figure S3D). The geometry of the microperifusion platform was modeled as a 2D cross-section of the microfluidic device with fluid flow from left to right (Figure S3E). In both the macroperifusion and microperifusion models, 5 islets with a diameter of 150 µm (5 IEQs) were placed in the flow path. FEM models were solved as a time-dependent problem, allowing for intermediate time-steps that corresponding with fraction collection time during macro and microperifusion. A list of the parameters used in the computational models is provided (Table S3).

### Statistics

Data were expressed as mean ± standard error of mean. A p-value less than 0.05 was considered significant. Analyses of area under the curve and statistical comparisons (Mann-Whitney test, Wilcoxon matched-pairs signed rank test, and one- and two-way ANOVA) were performed using Prism v8 software (GraphPad, San Diego, CA). Statistical details of experiments are described in the Figure Legends and Results.

