## Supplemental Information for "Human pseudoislet system enables detection of differences in G-protein-coupled-receptor signaling pathways between α and β cells"

### Figure S1

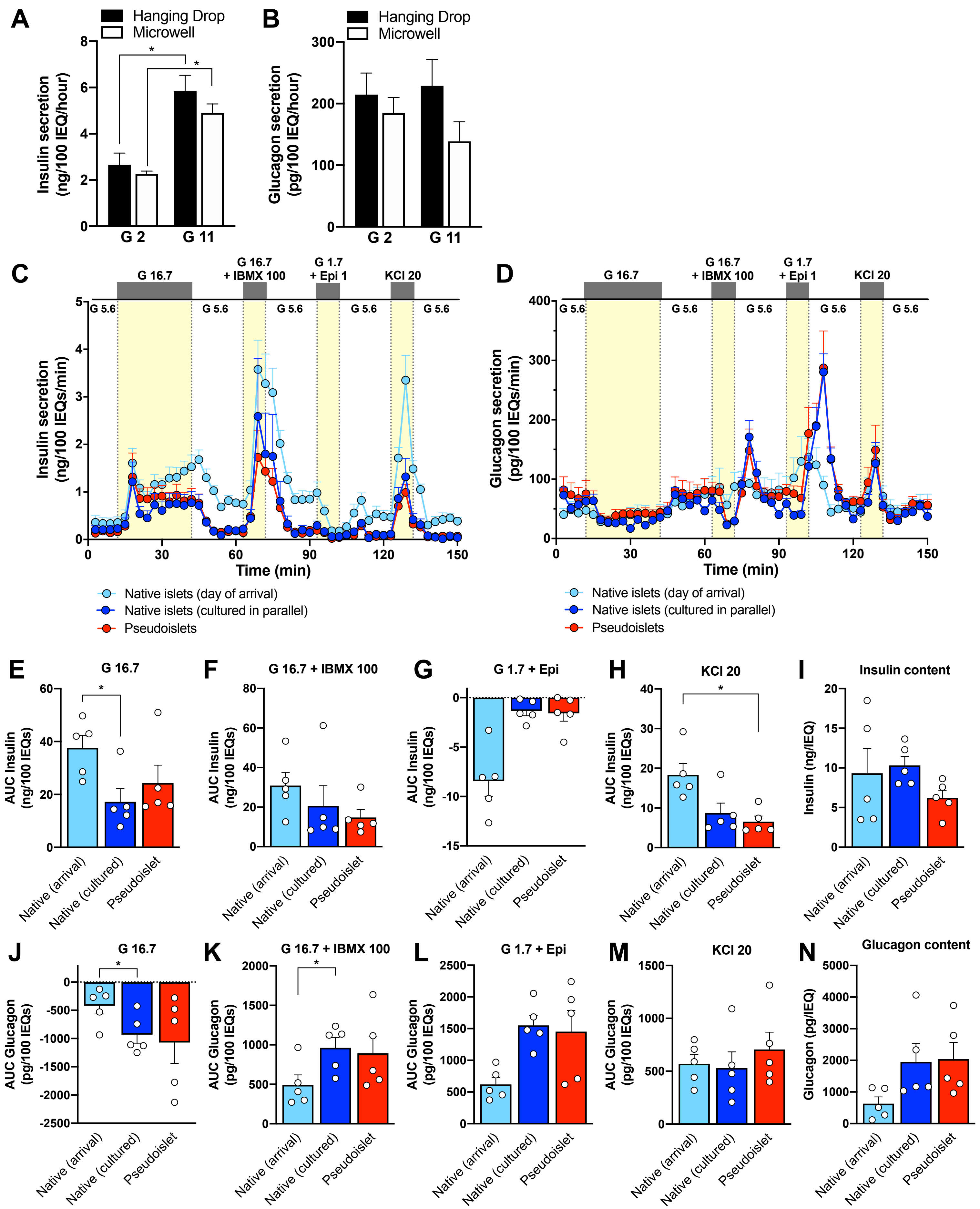

A
Hanging Drop
Microwell
Insulin secretion (ng/100 IEQ/hour)
G 2
G 11
B
Glucagon secretion (pg/100 IEQ/hour)
C
G 16.7
G 16.7 + IBMX 100
G 1.7 + Epi 1
KCl 20
G 5.6
Insulin secretion (ng/100 IEQs/min)
Time (min)
Native islets (day of arrival)
Native islets (cultured in parallel)
Pseudoislets
D
Glucagon secretion (pg/100 IEQs/min)
E
G 16.7
AUC Insulin (ng/100 IEQs)
Native (arrival)
Native (cultured)
Pseudoislet
F
G 16.7 + IBMX 100
G
G 1.7 + Epi
H
KCl 20
I
Insulin content
Insulin (ng/IEQ)
J
AUC Glucagon (pg/100 IEQs)
K
L
M
N
Glucagon content
Glucagon (pg/IEQ)

**Figure S2**

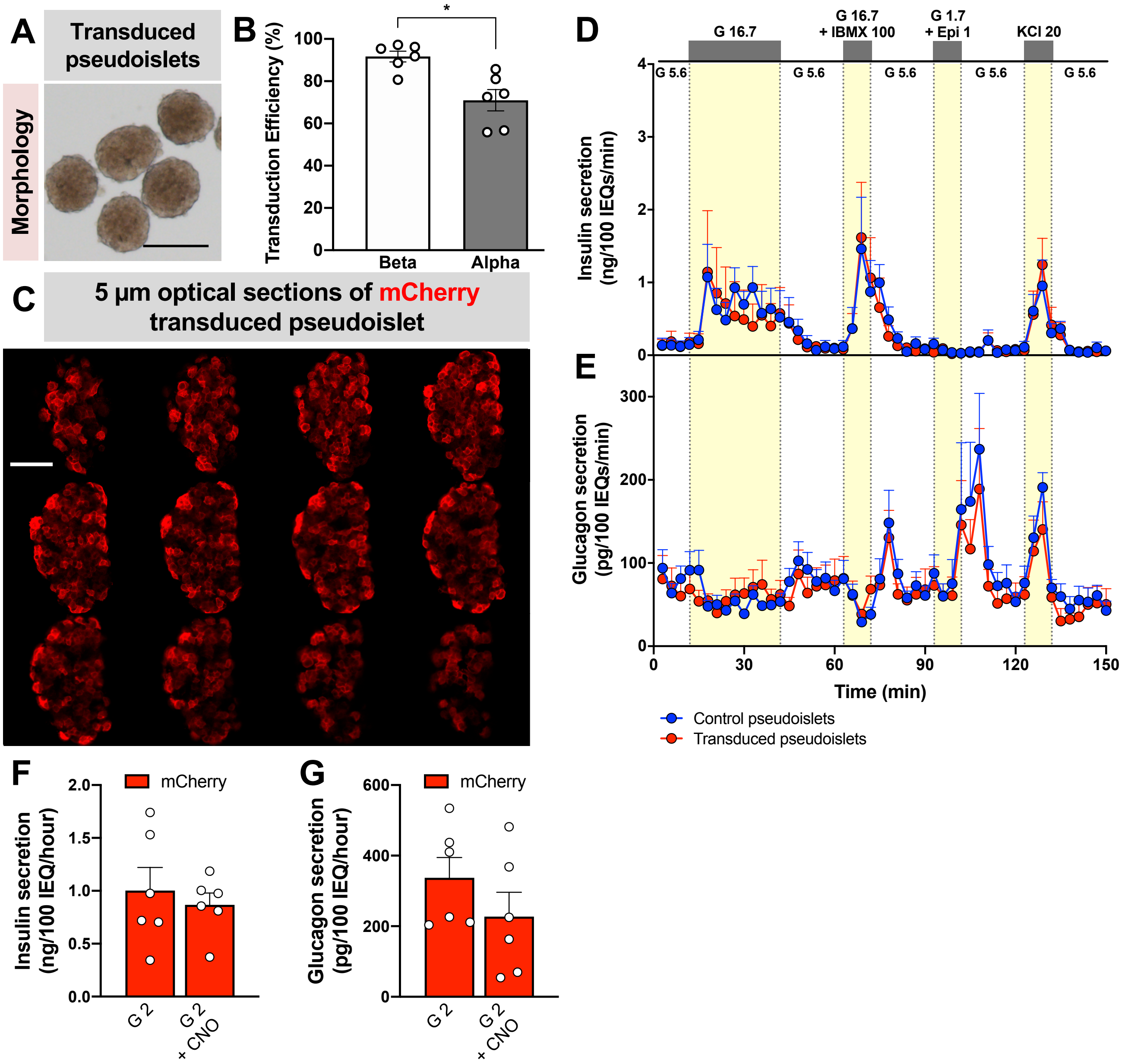
A
Transduced pseudoislets
Morphology
B
Transduction Efficiency (%)
Beta
Alpha
C
5 µm optical sections of mCherry transduced pseudoislet
D
G 16.7
G 16.7 + IBMX 100
G 1.7 + Epi 1
KCl 20
G 5.6
Insulin secretion (ng/100 IEQs/min)
E
Glucagon secretion (pg/100 IEQs/min)
Time (min)
Control pseudoislets
Transduced pseudoislets
F
mCherry
Insulin secretion (ng/100 IEQ/hour)
G 2
G 2 + CNO
G
mCherry
Glucagon secretion (pg/100 IEQ/hour)
G 2
G 2 + CNO

**Figure S3**

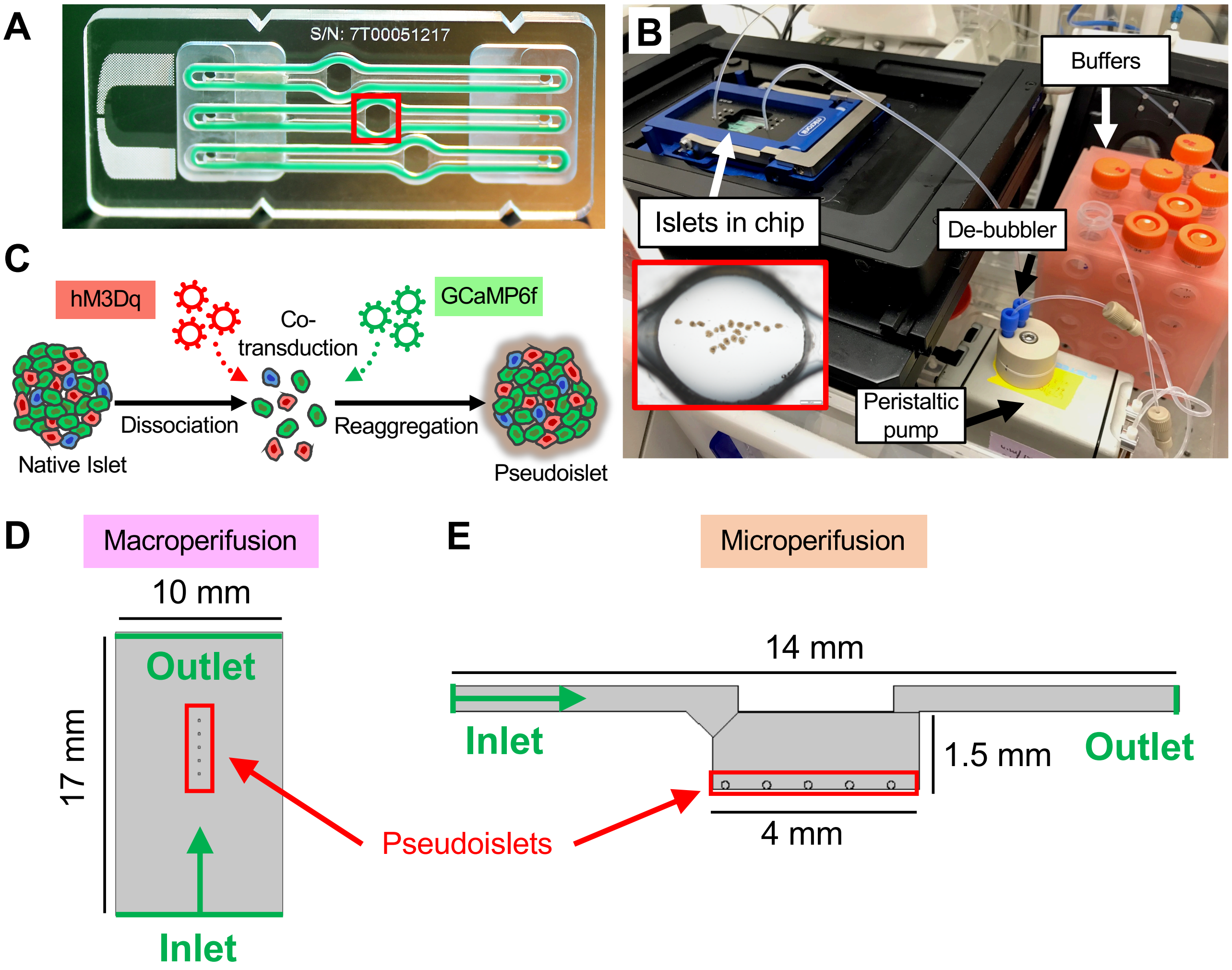


**F**

| Perifusion parameters | Macro | Micro |
|---|---|---|
| Chamber volume (μL) | 1500 | 65 |
| Flow rate (μL/min) | 3000 | 100 |
| Fraction time (mins) | 3 | 2 |
| Stimulus time (mins) | 21 | 8 |
| Islet equivalents | ~250 | ~25 |

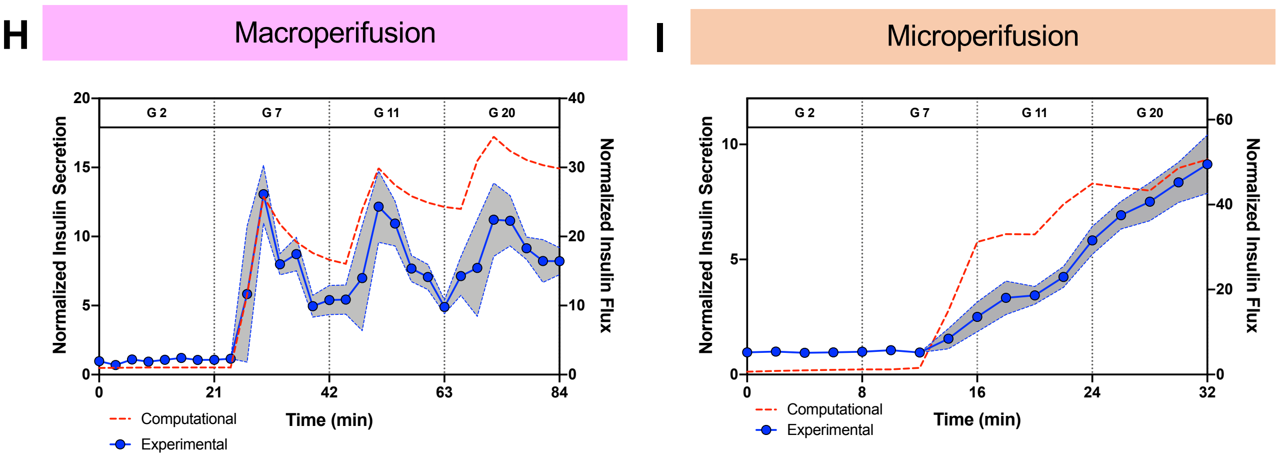

#### SUPPLEMENTAL FIGURE LEGENDS

**Figure S1. Related to Figure 1. Evaluation of hormone secretory responses in pseudoislet system.** Comparison of insulin (A) and glucagon secretion (B) by static incubation in pseudoislets made via modified hanging drop system (InSphero) versus ultra-low attachment microwell system (Perkin Elmer); n=1 donor, 3 replicates, $p > 0.05$. Insulin (C) and glucagon secretion (D) was measured by macroperifusion in native islets on the day of arrival to Vanderbilt (light blue trace) was compared in the same donor with secretory response of native islets cultured for six days in Vanderbilt pseudoislet media (dark blue trace, also shown in Figures 1H and 1I) and pseudoislets collected at the end of six day reaggregation period (red trace, also shown in Figures 1H and 1I); n=5 donors; G 5.6 – 5.6 mM glucose; G 16.7 – 16.7 mM glucose; G 16.7 + IBMX 100 – 16.7 mM glucose with 100 µM isobutylmethylxanthine (IBMX); G1.7 + Epi 1 – 1.7 mM glucose and 1 µM epinephrine (Epi); KCl 20 – 20 mM of potassium chloride (KCl). The area under the curve (AUC) of the insulin secretory responses to G 16.7 (E), G 16.7 + IBMX 100 (F), G 1.7 +Epi 1 (G), KCl 20 (H) and insulin content (E). The AUC of the glucagon secretory responses to G 16.7 (J), G 16.7 + IBMX 100 (K), G1.7 + Epi 1 (L), and KCl 20 (M), and glucagon content (N). One-way ANOVA with Dunn's multiple comparison test was used to analyze differences in panels E-N; *, $p<0.05$.

**Figure S2. Related to Figure 3. Pseudoislet system allows for highly efficient transduction of human islet cells.** (A) Bright-field image of transduced pseudoislets. Scale bar is 200 µm. (B) Transduction efficiency of β and α cells in this system; *, $p < 0.05$. (C) Confocal image of mCherry-transduced pseudoislet showing optical sections taken every 5-µm to highlight transduced cells throughout the entire pseudoislet. Scale bar is 100 µm. Insulin (D) and glucagon (E) secretion in response to a series of β and α cell secretagogues measured by macroperifusion in control pseudoislets and pseudoislets transduced with m-Cherry virus from the same donor (n=3 donors). Insulin (F) and glucagon secretion (G) in mCherry-expressing pseudoislets measured in 2 mM glucose (G 2) with and without CNO (n=2 donors; n=3 replicates/donor; $p > 0.05$). Panels D and E were analyzed by 2-way ANOVA; $p > 0.05$. Panels F and G were compared by Mann-Whitney test.

**Figure S3. Related to Figure 5. Microperifusion system assembly and fluid dynamic modeling of macro- and microperifusion.** (A) Picture of microfluidic device showing where islets are loaded and imaged (red box). There are three potential chambers to load islets, but in the current experimental layout, islets are loaded only into one chamber. (B) Experimental set-up of the microfluidic device on the confocal microscope stage within incubator including peristaltic pump, de-bubbler, and perifusion buffers. (C) Schematic of experimental workflow with incorporation of genetically encoded biosensor into hM3Dq-expressing pseudoislets. Schematic of the macroperifusion (D) and microperifusion chamber (E) showing the path of fluid flow. (F) Key experimental parameters of macroperifusion and microperifusion system. (H and I) Comparison of normalized insulin secretion acquired experimentally versus predicted by modeling in macroperifusion (H) and microperifusion (I). Experimental insulin data was normalized to average value in 2 mM glucose. The gray region demonstrates the SEM comparing experimental insulin secretion data and insulin flux from COMSOL computational modeling in the macroperifusion system (H) and microperifusion system (I); G 2 – 2 mM, G 7 – 7 mM, G 1 – 11 mM, G 20 – 20 mM glucose.

**Checklist for reporting human islet preparations used in research**
Adapted from Hart NJ, Powers AC (2019) Progress, challenges, and suggestions for using human islets to understand islet biology and human diabetes. Diabetologia https://doi.org/10.1007/s00125-018-4772-2 and Poitout V, Satin LS, Kahn SE, et al (2019) A call for improved reporting of human islet characteristics in research articles. Diabetologia and Diabetes. https://doi.org/10.2337/dbi18-0055

**Table S1. Human Islet Donor Information**

| Islet preparation | 1 | 2 | 3 | 4 | 5 | 6 | 7 | 8 |
|---|---|---|---|---|---|---|---|---|
| Unique identifier | DON308 | 08619527 | 08617638 | 10861888 | 11046361 | 11106978 | 11578698 | 12713942 |
| Donor age (years) | 35 | 28 | 49 | 58 | 57 | 20 | 57 | 41 |
| Donor sex | M | M | M | F | M | F | M | M |
| Donor BMI ($kg/m^2$) | 24.7 | 34.7 | 34.0 | 36.6 | 35.9 | 36.9 | 25.8 | 23.5 |
| Donor HbA1C | 5.3 | N/A | 6.1 | 5.3 | 5.4 | 5.3 | 5.7 | 5.2 |
| Origin/source of islets | AHN | IIDP | IIDP | IIDP | IIDP | IIDP | IIDP | IIDP |
| Islet isolation centre | AHN | SL | SC | SC | SL | SL | SL | SL |
| Donor history of diabetes? | No | No | No | No | No | No | No | No |
| Donor cause of death | Head Trauma | Head Trauma | Stroke | Anoxia | Stroke | Head Trauma | Stroke | Stroke |
| Warm ischaemia time (h) | N/A | N/A | N/A | 0.3 | N/A | N/A | N/A | N/A |
| Cold ischaemia time (h) | N/A | 14.2 | 8.5 | 6.4 | 9.2 | 6.5 | 6.8 | 9.8 |
| Estimated purity (%) | 50 | 95 | 80 | 80 | 80 | 83 | 95 | 85 |
| Estimated viability (%) | 65 | 95 | 97 | 97 | 95 | 95 | 95 | 95 |
| Total culture time (h) | 80 | 103 | 47 | 96 | 120 | 92 | 114 | 81 |
| Glucose-stimulated insulin secretion or other functional measurement | perifusion | perifusion | perifusion | perifusion | perifusion | perifusion | perifusion | perifusion |
| Handpicked to purity? | Yes | Yes | Yes | Yes | Yes | Yes | Yes | Yes |

| Islet preparation | 9 | 10 | 11 | 12 | 13 | 14 | 15 | 16 |
|---|---|---|---|---|---|---|---|---|
| Unique identifier | 10490796 | 10516338 | R252 | R253 | R260 | R264 | R282 | R286 |
| Donor age (years) | 53 | 52 | 26 | 57 | 73 | 44 | 57 | 41 |
| Donor sex | F | F | F | M | F | M | M | M |
| Donor BMI ($kg/m^2$) | 26.3 | 26.9 | 25.4 | 25.5 | 26.9 | 33.7 | 26.4 | 20.3 |
| Donor HbA1C | 5.4 | 4.5 | 5.0 | 5.0 | 6.2 | 5.7 | 6.0 | 5.2 |
| Origin/source of islets | IIDP | IIDP | ADI Islet Core | ADI Islet Core | ADI Islet Core | ADI Islet Core | ADI Islet Core | ADI Islet Core |
| Islet isolation centre | SL | SL | ADI Islet Core | ADI Islet Core | ADI Islet Core | ADI Islet Core | ADI Islet Core | ADI Islet Core |
| Donor history of diabetes? | No | No | No | No | No | No | No | No |
| Donor cause of death | Anoxia | Stroke | N/A | N/A | N/A | N/A | N/A | N/A |
| Warm ischaemia time (h) | N/A | N/A | N/A | N/A | N/A | N/A | N/A | N/A |
| Cold ischaemia time (h) | 7.6 | 6.3 | 11 | 14.3 | 11.5 | 11 | 9.5 | 13.5 |
| Estimated purity (%) | 90 | 80 | 85 | 90 | 80 | 90 | 90 | 95 |
| Estimated viability (%) | 95 | 95 | N/A | N/A | N/A | N/A | N/A | N/A |
| Total culture time (h) | 88 | 108 | 55 | 88 | 70 | 98 | 96 | 44 |
| Glucose-stimulated insulin secretion or other functional measurement | perifusion | perifusion | perifusion | perifusion | perifusion | perifusion | perifusion | perifusion |
| Handpicked to purity? | Yes | Yes | Yes | Yes | Yes | Yes | Yes | Yes |

| Islet preparation | 17 | 18 | 19 | 20 | 21 | 22 | 23 | 24 |
|---|---|---|---|---|---|---|---|---|
| Unique identifier | R306 | R309 | R314 | R318 | R323 | R340 | HPAP022 | HPAP035 |
| Donor age (years) | 22 | 47 | 31 | 54 | 60 | 36 | 39 | 35 |
| Donor sex | F | F | F | M | F | M | F | M |
| Donor BMI (kg/m$^2$) | 21.1 | 27.4 | 30.3 | 20.5 | 24.9 | 23.3 | 34.76 | 26.91 |
| Donor HbA1C | 5.3 | 5.5 | 5.0 | 5.0 | 5.1 | 5.3 | 4.7 | 5.2 |
| Origin/source of islets | ADI Islet Core | ADI Islet Core | ADI Islet Core | ADI Islet Core | ADI Islet Core | ADI Islet Core | HPAP | HPAP |
| Islet isolation centre | ADI Islet Core | ADI Islet Core | ADI Islet Core | ADI Islet Core | ADI Islet Core | ADI Islet Core | UPenn | UPenn |
| Donor history of diabetes? | No | No | No | No | No | No | No | No |
| Donor cause of death | N/A | N/A | N/A | N/A | N/A | N/A | Anoxia | Anoxia |
| Warm ischaemia time (h) | N/A | N/A | N/A | N/A | N/A | N/A | N/A | N/A |
| Cold ischaemia time (h) | 15 | 11 | 14.75 | 16 | 14.5 | 12.5 | 8.55 | 12.9 |
| Estimated purity (%) | 80 | 90 | 80 | 90 | 90 | 95 | 95 | 90 |
| Estimated viability (%) | N/A | N/A | N/A | N/A | N/A | N/A | N/A | N/A |
| Total culture time (h) | 55 | 108 | 72 | 85 | 90 | 65 | 72 | 55 |
| Glucose-stimulated insulin secretion or other functional measurement | perifusion | perifusion | perifusion | perifusion | perifusion | perifusion | perifusion | perifusion |
| Handpicked to purity? | Yes | Yes | Yes | Yes | Yes | Yes | Yes | Yes |

AHN – Allegheny Health Network, IIDP – Integrated islet distribution program, SL – The Scharp-Lacy Research Institute. SC – Southern California Islet Cell Resource Center, ADI IsletCore – Alberta Diabetes Institute IsletCore, HPAP – Human Pancreas Analysis Program,

**Table S2. Antibody Information**

| Antigen | Host Species | Dilution | Vendor/Source | Catalog # |
|---|---|---|---|---|
| **ARX** | Sheep | 1:1000 | R&D Systems | AF7068 |
| **C-peptide** | Rat | 1:1000 | DSHB | GN-ID4 |
| **Caveolin-1** | Rabbit | 1:2000 | Abcam | ab2910 |
| **Collagen-IV** | Rabbit | 1:1000 | Rockland | 600-401-106S |
| **GFP** | Chicken | 1:1000 | Abcam | ab13970 |
| **Glucagon** | Rabbit | 1:200 | Cell Signaling | 2760 |
| **Glucagon** | Mouse | 1:500 | Abcam | ab10988 |
| **Insulin** | Guinea pig | 1:1000 | Dako | A0564 |
| **Ki67** | Rabbit | 1:5000 | Abcam | ab15580 |
| **MAFB** | Rabbit | 1:3000 | Roland Stein | N/A |
| **mCherry** | Rabbit | 1:1000 | Abcam | ab167453 |
| **NKX2.2** | Mouse | 1:1000 | DSHB | 74-5A5 |
| **NKX6.1** | Rabbit | 1:2000 | BCBC/Palle Serup | N/A |
| **PAX6** | Rabbit | 1:5000 | Covance | PRB-28P-100 |
| **PDX1** | Rabbit | 1:5000 | C. V. E. Wright | N/A |
| **Somatostatin** | Goat | 1:500 | Santa Cruz | sc-7819 |
| **VEGFR2** | Goat | 1:200 | R&D Systems | AF644 |

BCBC – Beta Cell Biology Consortium, DHSB – Developmental Studies Hybridoma Bank, N/A – not applicable

#### Table S3. Computational Modeling Parameters

| | Parameter | Macroperifusion | Microperifusion |
|---|---|---|---|
| **Fluid Dynamics** | Fluid | Water | |
| | Density | 993 kg/$m^3$ | |
| | Dynamic Viscosity | $7x10^{-4}$ Pa/s | |
| | Inlet Velocity | $2.13x10^{-4}$ m/s | $3.34x10^{-2}$ m/s |
| **Mass Transport** | Diffusion Coefficient, Glucose in Fluid | $9x10^{-10}$ $m^2$/s | |
| | Diffusion Coefficient, Glucose in Islets | $3x10^{-10}$ $m^2$/s | |
| | Diffusion Coefficient, Oxygen in Fluid | $3x10^{-9}$ $m^2$/s | |
| | Diffusion Coefficient, Oxygen in Islets | $2x10^{-9}$ $m^2$/s | |
| | Diffusion Coefficient, Insulin in Fluid | $1.5x10^{-10}$ $m^2$/s | |
| | Diffusion Coefficient, Insulin in Islets | $0.5x10^{-10}$ $m^2$/s | |
| | Oxygen Concentration | 0.2 mol/$m^3$ | |
| **Islet Physiology** | Number of Islets | 5 | |
| | Radius | $7.5x10^{-5}$ m | |
| | Maximum Oxygen Consumption Rate | -0.034 mol/s/$m^3$ | |
| | Maximum Glucose Consumption Rate | -0.028 mol/s/$m^3$ | |
| | Insulin Release Rate Constant | $3x10^{-3}$ 1/s | |
| | Maximum First Phase Insulin Secretion Rate | $10x10^{-5}$ mol/s/$m^3$ | |
| | Maximum Second Phase Insulin Secretion Rate | $1.8x10^{-5}$ mol/s/$m^3$ | |

*Parameters in center apply to both perifusion systems
